# Single-cell mRNA profiling reveals heterogeneous combinatorial expression of *Hoxd* genes during limb development

**DOI:** 10.1101/327619

**Authors:** P. J. Fabre, M. Leleu, B. Mascrez, Q. Lo Giudice, J. Cobb, D. Duboule

## Abstract

A global analysis of gene expression during development reveals specific transcription patterns associated with the emergence of various cell types, tissues and organs. These heterogeneous patterns are instrumental to ensure the proper formation of the different parts of our body, as shown by the phenotypic effects generated by functional genetic approaches. However, variations at the cellular level can be observed within each structure or organ. In the developing mammalian limbs, expression of *Hoxd* genes is differentially controlled in space and time in cells that will pattern the digits and the arms. Here we analyze single-cell transcriptomes of limb bud cells and show that *Hox* genes are expressed in specific combinations that match particular cell types. In the presumptive digits, we find that the expression of *Hoxd* gene is unbalanced, despite their common genomic proximity to known global enhancers, often expressing only a subset of the five genes transcribed in these cells. We also report that combinatorial expression follows a pseudo-time sequence, suggesting that a progression in combinatorial expression may be associated with cellular diversity in developing digits.

**HIGHLIGHTS:**

- Collinear expression of Hox genes is only weaved at the tissue scale
- Enhancer-sharing to specific target genes is reduced at the single-cell level
- Hoxd gene combinatorial expression is linked to distinct transcriptional signatures
- In presumptive digits, Hoxd combinations follow a pseudotime trajectory

## INTRODUCTION

Limb morphogenesis is controlled by several key transcription factors, amongst them members of the *Hox* gene family, in particular genes from the *HoxA* and *HoxD* clusters. During early limb development, the posterior *Hoxd* genes are expressed in precise, partly overlapping domains ({Dolle, 1989a #13}{Dolle, 1993 #14}), which will pre-figure the various parts of the future appendices, i.e. the hands and feet (autopods) and the more proximally located arm (stylopod) and forearm (zeugopod) segments. Recently, it was shown that expression of *Hoxd9* to *Hoxd13* in presumptive digits is under the control of the same set of enhancer elements, located in the gene desert centromeric to the cluster itself {Montavon, 2011 #4}{Andrey, 2013 #2}{Fabre, 2017 #9}). However, their global expression patterns display some difference, with a broader expression of *Hoxd13* within future digit 1 (the thumb), whereas *Hoxd9* to *Hoxd12* transcripts were found only in future digits 2 to 5. This difference is likely due to the existence of a quantitative collinearity ({Dolle, 1991 #9} {Spitz, 2003 #16}{Montavon, 2008 #4}), whereby a gradual increase in the amount of steady-state mRNA levels is observed from *Hoxd9*, expressed at the weakest level, to the robust transcription of *Hoxd13*.

By using DNA FISH, the extent of chromatin interactions between *Hox* genes and their enhancers in single cells showed variability ({Fabre, 2017 #9}{Rodriguez-Carballo, 2017 #8}), and super-resolution microscopy confirmed that the *HoxD* gene cluster can display a variety of structural conformations in various future autopod cells ({Fabre, 2015 #2}). This heterogeneity is hard to reconcile with chromosome conformation datasets produced at this locus (e.g. {Montavon, 2011 #5} {Fabre, 2015 #32}), since the latter approach reflects the averaged behaviors of a cellular population. Consequently, a higher variability can be expected in cell-specific *Hox* gene transcriptions, when compared to previously reported expression profiles ({Duboule, 1989 #33}{Dolle, 1989 #15}({Montavon, 2008 #4}).

While genetic approaches have revealed the critical function of these genes during limb outgrowth and patterning, the homogeneous or heterogeneous impact of mutations at the cellular level is more difficult to evaluate. The ablation of *Hoxd13* alone leads to a morphological effect in digits weaker than when a simultaneous deletion of *Hoxd11, Hoxd12* and *Hoxd13* is achieved ({Dolle, 1989a #13} {Zakany, 1997a #12}{Delpretti, 2012 #17}), suggesting that *Hoxd11, Hoxd12* and *Hoxd13* functionally cooperate during digit development. However, how this cooperation occurs at the cellular level is unknown. This question is made even more complex by the regulatory strategies that evolved at the *HoxD* locus, where several neighbor genes can be regulated by several enhancers in the same large domains.

It was indeed recently reported that this cluster lies between two large topologically associating domain (TADs) ({Dixon, 2012 #34}{Andrey, 2013 #1}{Beccari, 2016 #35}{Fabre, 2015 #32}), each of them containing a range of enhancer elements acting in the same domains. The TAD located centromeric to *HoxD* (C-DOM) contains several enhancers specific for autopod (digit) cells, whereas T-DOM, the TAD located telomeric to *HoxD*, hosts a series of enhancers specific for future arm and forearm cells ({Andrey, 2013 #1}). In addition, genes with a central position in the cluster such as *Hoxd9, Hoxd10* or *Hoxd11* are targeted by enhancers belonging to the two different TADs, initially in zeugopod cells, then in autopod cells, suggesting an even greater heterogeneity in transcripts distribution. In order to try and evaluate *Hoxd* transcript heterogeneity during limb development, we produced single-limb cell transcriptomes of different origins, to see whether the apparently homogenous behavior in *Hox* gene transcriptional program as observed upon large-scale analyses was confirmed at the cellular level. We report here that *Hoxd* genes transcripts are present in various combinations in different limb cells and discuss the importance of these results in our understanding of how *Hoxd* genes are regulated and how their global functions are achieved in these structures.

## RESULTS

### Heterogeneity of posterior *Hoxd* gene transcripts in single-cells

In order to document the expression pattern of *Hoxd13* at the single-cell level, embryonic day (E) 12.5 limb sections were use in RNA-FISH experiments (**Fig. 1A**). As expected, we observed a high expression specificity in presumptive digits cells in the distal part of the forelimb, with the highest transcript levels in cells located at the boundary between the digital and the interdigital compartments, while lower levels were scored in interdigital mesenchyme. Signal was neither detected within the digital compartment, nor in more proximal parts of the limb ({Montavon, 2008 #4}) (**Fig. 1A**). However, a high heterogeneity in gene expression was recorded, with stippled signal pattern contrasting with the broader expression domain previously described. As a consequence, we asked whether all cells expressing *Hoxd13* would also contain *Hoxd11* transcripts, given that both genes are under the same regulatory control in these distal cells ({Montavon, 2011 #5}{Andrey, 2013 #1}). We micro-dissected autopod tissue to obtain a single-cell suspension and performed double fluorescent RNA labelling. The single-cell preparation was then analyzed by FACS and revealed that only a minority of cells were in fact expressing detectable levels of *Hoxd11* and/or *Hoxd13* (**Fig. 1B**). Amongst positive cells, the largest fraction was *Hoxd13* positive and negative for *Hoxd11* (*d13+d11-*; 53%), whereas double positive cells (*d13+d11+*) represented 38% only and 9% of the cells contained *Hoxd11* mRNAs alone (*d11+*)(**Fig. 1B**).

**Figure 1.**
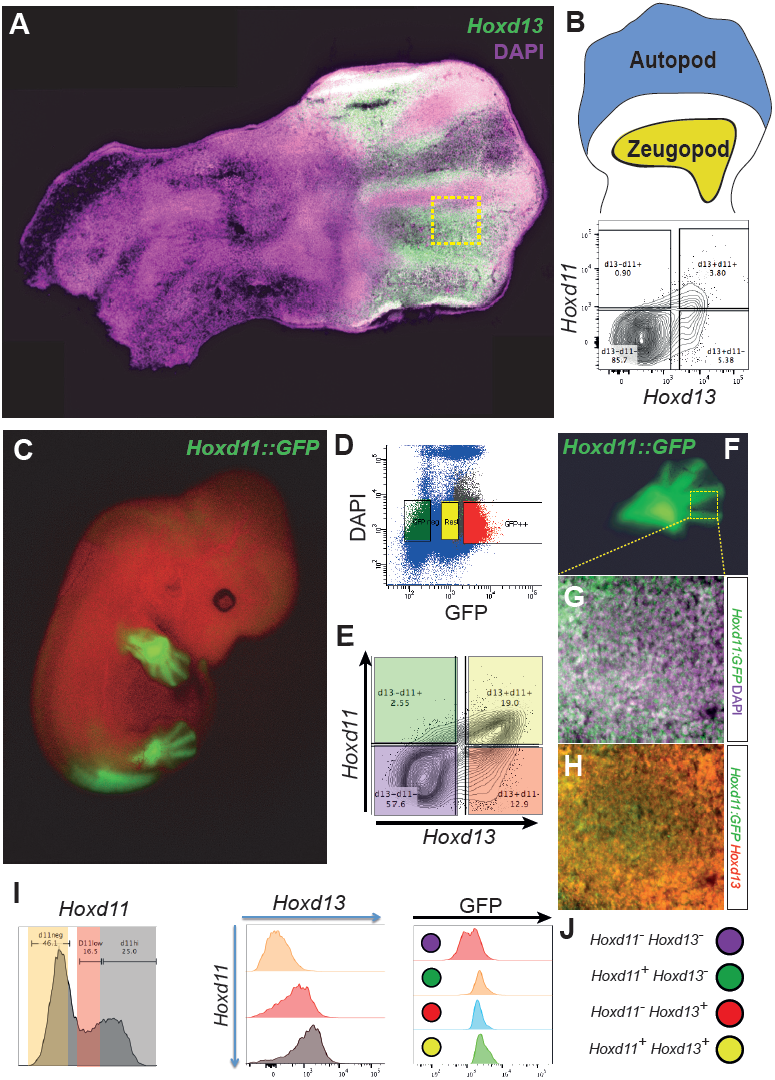
Heterogeneity of posterior *Hoxd* mRNAs in single-cells. **A**. Fluorescent *in situ* hybridization of *Hoxd13* mRNA on section reveals discrete expression pattern in the autopod of E12.5 mouse forelimb. **B**. Single-cell double-labelling of *Hoxd13* and *Hoxd11* mRNA from E12.5 autopod cells (up, schematic) by fluorescent hybridization followed by flow cytometry detection. The density plot (below) shows a high proportion of double negative cells. **C**. *Hoxd11* expression pattern revealed using a *Hoxd11::GFP* knock-in mouse strain, with high expression in digits and low expression in interdigital cells of autopod forelimbs, together with strong signals in zeugopod cells. **D-E**. Scatterplot profile from FACS to enrich for *Hoxd11-*positive cells using cells with high levels of GFP fluorescence (**D,** red) subsequently double-labelled for both *Hoxd13* and *Hoxd11* mRNAs. **E**. Flow cytometry analysis from the GFP-positive cells from **D** are shown in a density contour plots where colors highlight the four population of cells expressing various levels of *Hoxd11* and *Hoxd13*. **F-H**. Double-labeling of GFP (green, marker of *Hoxd11*-positive cells, **F-G**) and *Hoxd13* (**H**, red, FISH) with DAPI (magenta) suggests four different combinations of *Hoxd* positive cells: Double positive for *Hoxd13* and *Hoxd11*, single *Hoxd13-*positive; single *Hoxd11*-positive and double negative for *Hoxd13* and *Hoxd11.* **I.** Histograms showing the correlation of expression where high levels of *Hoxd13* are associated with high level of *Hoxd11* (two panels on the left). Higher levels of GFP are also observed in cells expressing at least one of the *Hoxd* genes (right panel). **J.** Schematic of the four different combinations observed in **E** and **G-H**: double negative for *Hoxd13* and *Hoxd11* (purple); single *Hoxd11*-positive (green), single *Hoxd13-*positive (red) and double positive *Hoxd13* and *Hoxd11* (yellow).

Because a substantial number of cells did not express any *Hoxd* genes, we enriched for the positive fraction using a mouse line containing a GFP reporter sequence knocked in *Hoxd11*. In these mice, GFP was produced in those cells where *Hoxd11* had been transcribed (**Fig. S1**). We monitored the fluorescence at E12.5 and observed a pattern recapitulating *Hoxd11* endogenous expression ({Dolle, 1989a #15; Dolle, 1989b #13}) (**Fig. 1C**). E12.5 limb cells from these animals were FACS sorted using the GFP (**Fig. 1D**) and, under these conditions, the double labelling of GFP positive cells increased to more than a third of the cells (**Fig. 1E**). However, amongst the positive cells, the ratio between the three *Hoxd* positive populations (*Hoxd13* only, *Hoxd11* only and double-positive) was roughly the same as before (37%, 7% and 55%, respectively). To confirm the presence of these different populations, we performed *Hoxd13* RNA-FISH on sections from *Hoxd11::GFP* E12.5 forelimbs (**Fig. 1F**) and observed a high variability in GFP levels (**Fig. 1G**). We found that high levels of *Hoxd13* were observed in cells with either little or no *Hoxd11* activity (**Fig. 1H**), yet the majority of cells displayed high signals for both *Hoxd11* and *Hoxd13*, suggesting that in these cells the two genes were regulated in a similar manner.

To quantify a potential correlation between *Hoxd11* and *Hoxd13* expression levels in these GFP-positive cells, we binned *Hoxd11* positive cells in three categories: negative cells (*d11neg*, orange), cells expressing at low levels (*d11low*, red) and cells expressing at high levels (*d11hi*, grey; **Fig. 1I**, left panel). Flow cytometry analysis revealed that higher *Hoxd13* levels were clearly observed in the *d11hi* population, indicating that in single cells, whenever both genes are expressed, they tend to respond to enhancers with the same efficiency (**Fig. 1I**). To relate these latter results with the level of GFP observed by microscopy (**Fig. 1 H**), we monitored the levels of GFP in single-cells and found a correlation between abundant *Hoxd11* mRNAs and higher levels of the GFP protein (**Fig. 1I**, right panel). Altogether, these results suggested that some cellular heterogeneity exists with respect to *Hoxd* gene transcription in presumptive digit cells, with the possibility for sub-populations of cells to selectively express either one or the two genes. Overall, these observations contrast with the view that all limb cells transcribe all posterior *Hoxd* genes, a view conveyed by the global analysis of expression patterns by whole-mount *in situ* hybridization (WISH).

### Single limb cells transcriptomics

To obtain a wider view of this cellular heterogeneity as well as to see whether it depends on the position and fate of various limb cells, we performed single-cell RNA-seq to expand the analysis to all *Hox* genes. Because of its potential to detect as little as single-digit input spike-in molecules, we used the Fluidigm microfluidics C1 captures to obtain the maximal intensity of transcript detection ({Svensson, 2017 #36}). To enrich for cells expressing at least one *Hoxd* gene, we used only the GFP positive cells sorted by flow cytometry from the *Hoxd1I::GFP* mouse E12.5 forelimbs (see **Fig. 1C-E**). After capture, the cells were sequenced at very high depth to reach the finest sensitivity of gene detection, with an average of about 8.7 M reads per cell (**Fig. S2**).

We first showed that autopod and zeugopod cells portray distinct transcriptional signatures that can be observed in a machine learning algorithm that reduces dimensionality (tSNE). In this plot representation, we saw only little intermingling between autopod and zeugopod cells (**Fig. 2A-B**). To ensure that the single-cells signatures were specific to the two populations, we performed a differential expression analysis between the distal and proximal limbs. As shown in the MA-plot we found that genes specific to one or the other populations were indeed known markers of the two tissues (**Fig. 2C**). In fact, most of the autopod-specific genes are part of a tight interactive network established through weighted aggregation of known interactions (**Figs. 2D** and **S3**), thus demonstrating the high level of gene detection in our single-cells (**Figs. 2D** and **S2**).

**Figure 2.**
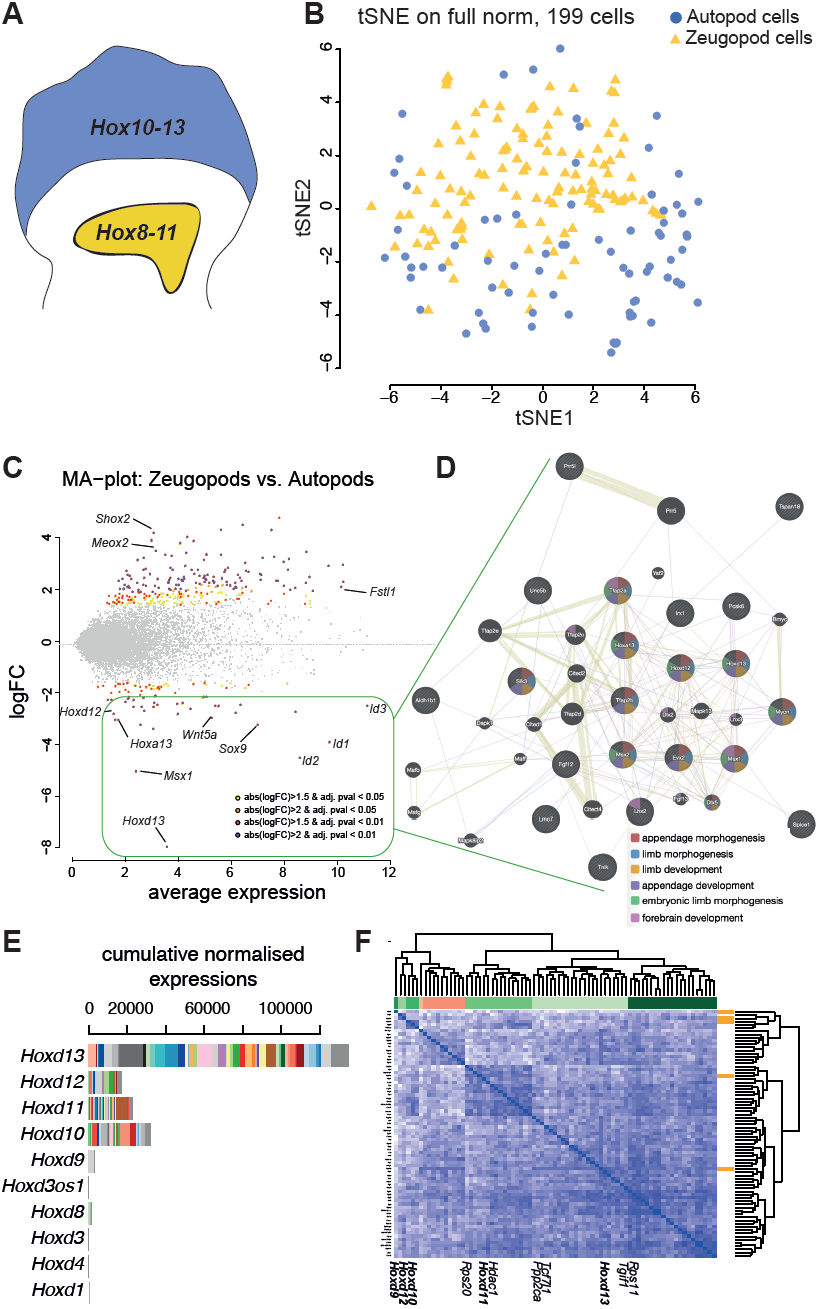
Single-cell transcriptomics from forelimbs autopod and zeugopod. **A.** Schematics of forelimb territories harboring different combinations of *Hoxd* genes activity. Single-cell RNA-seq was performed on micro-dissected 12.5 forelimb autopods (AP, blue) and zeugopods (ZP, yellow) tissues derived from *Hoxd11::GFP* limbs and positive for GFP. **B.** T-distributed stochastic neighbor embedding (tSNE) plot of gene expression relationships amongst the 199 single-cells from AP (blue) and ZP (yellow) shows a segregation along the tSNE2 axis. **C.** MA-plot produced from cross-analysis between AP and ZP cells. Known genes with differential expression between tissues are indicated on the graph. **D.** Gene nodes from the differentially expressed genes (DEG) up-regulated in autopod tissue, as computed using co-expression and interaction meta-analysis. **E.** Cumulative combinatorial expression of *Hoxd* genes in the autopod cells indicating variability of gene expression across *Hoxd* genes from cell to cell. **F.** Spearman’s rank correlation heatmap and hierarchical clustering of genes that covaried with at least one posterior *Hoxd* gene in the autopod cells. Below are shown the names of *Hoxd* genes (bold) and their known target genes.

To visualize the relative mRNA contributions of all *Hoxd* genes, we plotted their cumulative expressions with color coded single-cell (**Figs. 2E and S4**). While the distribution of absolute levels mirrored quite well the pattern previously established using other approaches ({Fabre, 2017 #9; Montavon, 2008 #4; Montavon, 2011 #5}), we observed again a selectivity of expression, which also applied to *Hoxd12, Hoxd11* and *Hoxd10*. Of note, amongst autopod cells positive either for *Hoxd13* and/or for *Hoxd11*, we identified similar proportions as before, with the majority of cells expressing both *Hoxd11* and *Hoxd13*, 40 percent containing *Hoxd13* mRNAs only and 10 percent with only *Hoxd11* mRNAs. To assess the potential covariances between the five posterior *Hoxd* genes (from *Hoxd9* to *Hoxd13*), we classified by Spearman’s rank correlation the genes that covaried with at least one of the *Hoxd* genes. A hierarchical clustering from these 76 genes showed a clear segregation between *Hoxd11/Hoxd13* on the one hand, and *Hoxd9, Hoxd10 and Hoxd12*, on the other hand (**Fig. 2F**). While *Hoxd9, Hoxd10* and *Hoxd12* were closely associated in the presence or absence of their mRNAs, *Hoxd11* and *Hoxd13* were part of two different sub-clusters associated with different set of genes, suggesting that the cell-specific expression of combinations of *Hoxd* genes may have biological relevance.

### Combinatorial *Hoxd* genes expression observed in single-cells

Therefore, despite their shared tissue-specific regulatory landscapes, all *Hoxd* genes are not systematically expressed by the same cells. A discretization of the expression levels allowed us to score the various mRNA combinations observed either in autopod (**Fig. 3A**), or in zeugopod (**Fig. S5**) single-cells. In the autopod, the largest population was composed of cells expressing *Hoxd13* only, followed by a population expressing both *Hoxd11* and *Hoxd13* and then by an unexpected pool of cells with only *Hoxd10* and *Hoxd13* mRNAs. Cells containing three or more distinct *Hoxd* mRNAs were a minority and only 11 percent of cells expressed four genes, from *Hoxd10 to Hoxd13*. We asked whether these unambiguous associations were random or coupled with specific gene signatures by performing a tSNE on all autopod and zeugopod cells and we observed that groups of cells containing different combinations of *Hoxd* mRNAs tend to segregate, suggesting that their differences in gene expression is not restricted to *Hoxd* genes only (**Fig. 3C**).

**Figure 3.**
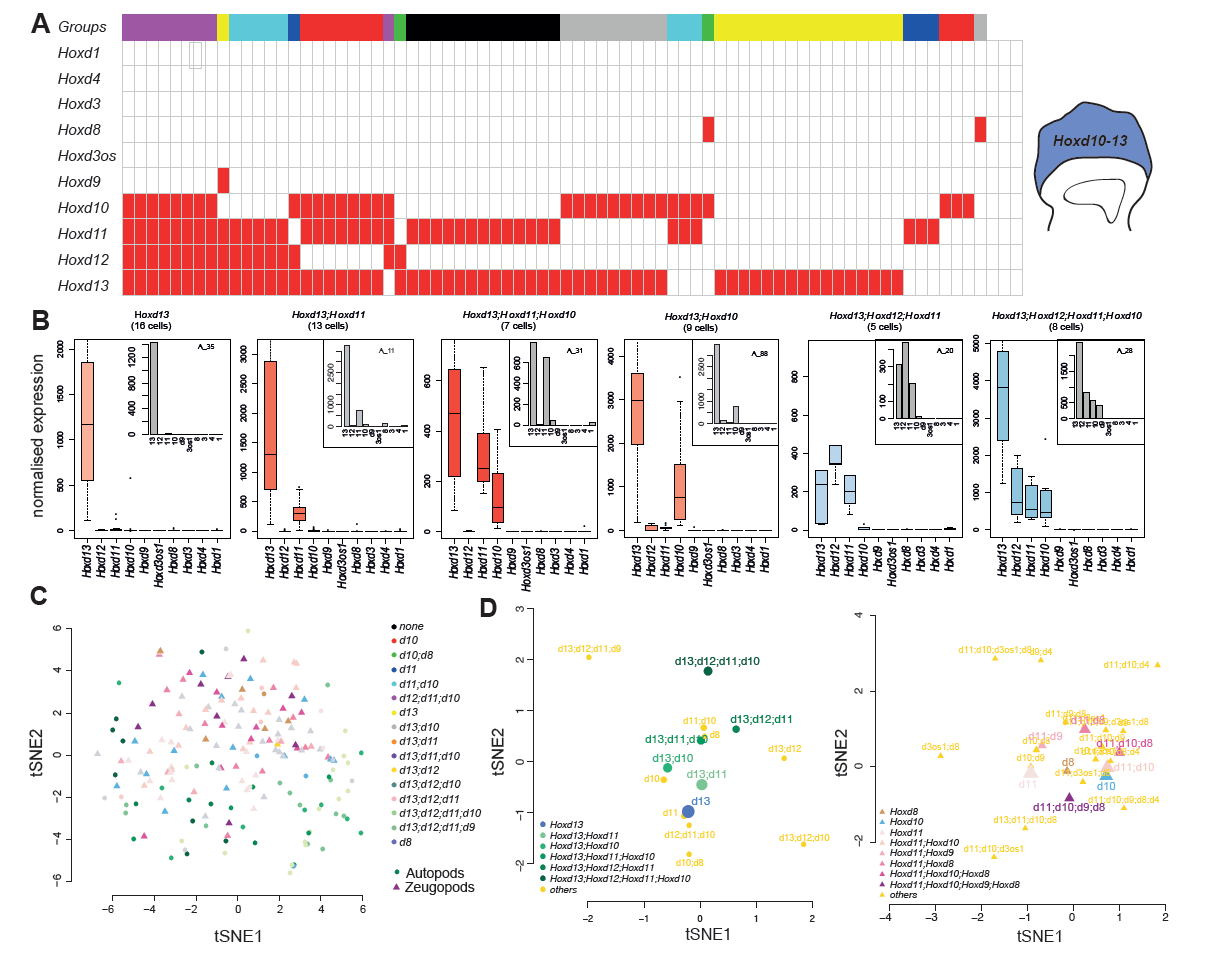
Combinatorial *Hoxd* genes expression in single-cells. **A**. Supervised cluster analysis reveals 16 combinations (clusters) of posterior *Hoxd* genes in autopod single-cells. **B**. Boxplots showing normalized expression for all *Hoxd* genes for the six main clusters from autopod cells, with a representative single cell shown in the top-right corner. tSNE with color-coded combinations of posterior *Hoxd* genes in autopods and zeugopods. tSNE showing groups of cells sharing the same combination of expression in autopod (left) and zeugopod (right).

We performed separate tSNE for autopod and zeugopod cells by clustering cells according to their *Hoxd* combinatorial patterns (**Fig. 3D**) and observed that some combinations tend to cluster together. This effect was particularly clear in autopod cells whenever a sufficient number of cells (>5) was plotted and we noticed that the transcriptional diversity increased along the second dimension of the tSNE when a higher diversity of *Hoxd* mRNAs was scored in the same cells. In zeugopod cells, groups of cells also segregated, though not as distinctly, suggesting a more homogeneous distribution of *Hoxd* mRNAs. These results suggested that subpopulation of autopod cells transcribe various combinations of *Hoxd* genes.

### Analysis of *Hoxd* cellular clusters

To more precisely assess this apparent cellular selectivity in *Hoxd* gene expression, we first determined whether particular cell clusters were at a specific phase of the cell cycle. While most cells with G2 scores were observed with either *Hoxd13* mRNAs only or with four posterior *Hoxd* genes active, we did not detect any significant difference associated with a specific combination of mRNAs (**Fig. S5**). We next performed a differential gene expression analysis to assess the degree of relationship between the six main cellular groups (**Fig. 4A-C**). Most of the differentially expressed genes (343 genes, **Table S1, and Fig. S6**) were scored between cells expressing only *Hoxd13* and cells expressing either three (*Hoxd11-Hoxd13*), or four genes (*Hoxd10* to *Hoxd13*) (**Fig. 4A**). Amongst these differentially expressed genes, many displayed strong autopod expression, including *Jag1*, which is downregulated in the absence of the HOX13 proteins ({Sheth, 2016 #18}. Out of 31 genes differentially expressed between cells containing either *Hoxd13* and *Hoxd11* mRNAs, or *Hoxd10, Hoxd11, Hoxd12* and *Hoxd13* mRNAs, only eight were specific to these two combinations (*Smarcc1, Mrps17, Snrpd2, Supt6, Tax1bp1, Rab5c, Ncbp2, Map3k7*).

**Figure 4.**
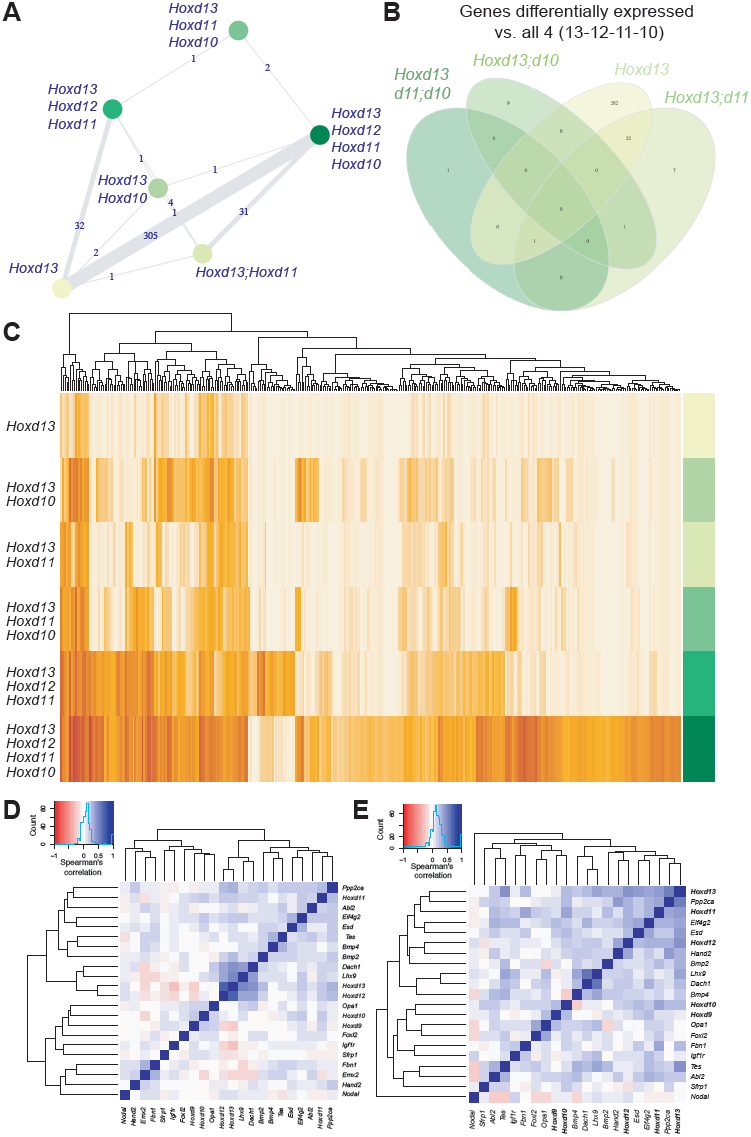
Analysis of *Hoxd* mRNAs combinations clusters. **A**. Network diagram of differentially expressed genes between the six main combinations of posterior *Hoxd* genes. **B**. Venn diagram showing the number of overlaping genes differentially expressed between the same combinations. **C**. Heatmap and unsupervised clustering of the 343 differentially expressed genes in the six groups of cells. **D-E**. Spearman’s rank correlation heatmaps and clusterings of posterior *Hoxd* genes and their targets in the full set of 199 cells from autopod and zeugopod (**D**) and only in autopod cells (**E**).

Noteworthy, clustering of expressed transcripts showed a hierarchical organization with a progression from those cells expressing *Hoxd13* only to two, then three and finally four *Hoxd* genes (**Fig. 4C**). As some of these genes were previously identified either as HOX proteins targets (e.g. *Ppp2ca {Salsi, 2008 #37}*), or being part of a *Hox* functional pathways (e.g. *Uty, Hoxa11os*), we assessed whether specific targets genes could be associated with particular combinations of *Hoxd* mRNAs. We generated a supervised clustering showing the covariance of known targets genes in a spearman correlation matrix (**Fig. 4D, E**). When the 199 cells originating from both the autopod and the zeugopod were considered, we found a clear partition of target gene mRNAs into two groups, corresponding to the nature of *Hoxd* mRNAs present (**Fig. 4D**). The presence of *Hoxd9* and *Hoxd10* mRNAs aggregated with targets genes such as *Hand2* and *Sfrp1*, whereas *Hoxd11, Hoxd12* and *Hoxd13* were co-expressed with different target genes such as *Ppp2ca* and *Bmp2/4*. Finally, the highest clustering across all cells was observed between *Hoxd12, Hoxd13, Dach1* and *Lhx9*, thus revealing a robust link between these genes.

When only autopod cells were considered, we observed two groups, with *Hoxd9* and *Hoxd10* transcripts in one cluster, while the more centromeric genes *Hoxd11, Hoxd12* and *Hoxd13* were transcribed in the other (**Fig. 4E**). As the former group did not express any of those genes typically up-regulated in distal cells (*Hoxd12* and *Hoxd13*), we wondered whether such differences in transcript distribution may reflect various stages in the progression of distal limb cells towards their final fates. Therefore, we implemented a measure of cellular pseudo-age, a strategy that evaluates a temporal hierarchy amongst single-cells based on their respective transcriptomes ({Haghverdi, 2016 #19}{Trapnell, 2014 #20}{Haghverdi, 2015 #49}). This approach allows to plot cells along a linearized axis to infer whether the combination alignments observed in the tSNE may correlate with a modulation of the time component.

We performed a pseudo-time analysis on the single-cells isolated from both the autopod and zeugopod and found that cells indeed spread along the pseudo-temporal axis that was linearized through a diffusion map (**Fig. 5A-B**). In these maps, while zeugopod cells did not distribute well along a temporal frame (**Fig. 5B**), the autopod cells were much better aligned (**Fig. 5A**). As demonstrated with gene expression clustering (**Fig 4. A-C**), specific combinations are distributed along the temporal axis in a way related to the various combinations of *Hoxd* mRNAs, with the *Hoxd13*-only cells at one extremity of the axis and the *Hoxd10 to Hoxd13* combination at the other extremity (**Fig. 5C-D**). Altogether, this clustering analysis showed that different combinations of *Hoxd* gene mRNAs may affect distinct groups of target genes. Noteworthy, it also revealed a preference for mRNA combinations involving neighbor genes, thus emphasizing the importance of genes’ position for their co-regulation.

**Figure 5.**
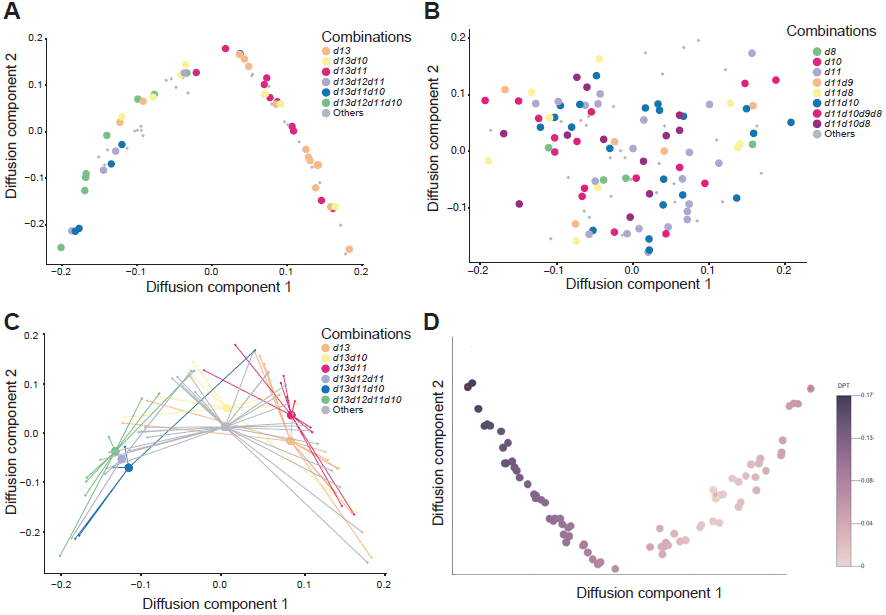
Diffusion pseudo-time across single-cells. **A.** Diffusion maps of the 76 autopod cells colored per *HoxD* combinations. This dot plot shows a progression from the simplest combinations to more complex ones. The most significant dimensions, i.e. the first two eigenvectors (DC1 and DC2) are displayed. **B.** Diffusion maps of the 120 zeugopod cells colored per *Hoxd* mRNA combinations. **C.** *Hoxd* groups centroid. The centroid of a *Hoxd* group, represented in the diffusion map of the 76 autopod cells. **D.** Distribution of autopod cells as shown in **A** are color coded to show their progression along diffusion pseudo-time.

## DISCUSSION

Limb bud cells require the expression of *Hox* genes originating from two separate clusters, *HoxA* and *HoxD*. We describe here the single-cell combinatorial expression of *Hoxd* genes found in cells sorted out by using a *Hoxd11*::GFP mouse strain. Albeit some cells tend to show higher level of *Hoxa* genes when the *Hoxd* genes were low, this was not the general rule. The fact that we did not score many *Hoxa*-mRNA positive cells after the enrichment for *Hoxd* genes expression (**Fig. S4**) may however reflect a compensatory mechanism whereby a strong global expression of one cluster would result in the weak transcription of the other. Distinct cellular content for either *Hoxd* or *Hoxa* mRNAs could account for the different phenotypic effects of inactivating these genes upon limb morphology, as exemplified by *Hoxd13* and *Hoxa13* ({Fromental-Ramain, 1996 #28}{Zakany, 1997 #30}{Kmita, 2005 #27};{Beccari, 2016 #35}). A more comprehensive single-limb cell sequencing strategy will fix this issue.

Our data show that *Hoxd* quantitative collinearity (Dollé et al., 1991) is to be considered at the global level since it results in fact from a sum of combinatorial expression of various genes in different cells. We emphasized that in autopod cells, the most frequently expressed gene is *Hoxd13*, as was expected from previous studies where it was described that this gene is expressed the strongest {Montavon, 2008 #4}. Apart from *Hoxd13*, other *Hoxd* genes were more sparsely activated, indicating either a stochastic process or a functional requirement for specific mRNA combinations in different cell types. This heterogeneous cellular situation raises two separate questions regarding first the underlying regulatory mechanism and, secondly, the potential functional significance.

### Different regulatory conformations?

We had previously shown that the regulation of posterior *Hoxd* genes in the distal limb bud was not entirely the same for all genes. In particular, *Hoxd13* is the only gene to be expressed in presumptive thumb cells, the other *Hoxd* mRNAs being excluding from this very digit ({Montavon, 2008 #4}). Also, the deletion of the *Hoxd13* locus lead to the upregulation of *Hoxd12* in thumb cells, yet not of the other remaining genes, suggesting that this thumb-specific expression was associated with the final and most 5’position of the gene on the cluster ({Kmita, 2002 #21}). The recent identification, in the posterior part of the *HoxD* cluster, of an unusually high density of bound CTCF molecules ({Soshnikova, 2010 #22}) may cause this transcriptional selectivity through the use of various sites, taking advantage of their distinct orientations ({Fabre, 2017 #9}{Rodriguez-Carballo, 2017 #8}).

In this view, the peculiar orientations of CTCF binding sites may allow the transient stabilization of various loop conformations, for example after extrusion driven by the cohesin complex ({Fudenberg, 2016 #44} {Haarhuis, 2017 #50}). Accordingly, distinct combinations of posterior *Hoxd* mRNAs could reflect the formation of specific loop extrusion patterns, in any single cell, as a choice between a fixed number of possibilities as determined by the presence of bound CTCF, with some conformations being favored over others. Of note we found that the cohesin-loading factor *Nipbl* is strongly downregulated in cells from the *Hoxd13* group when compared to cells expressing the full combination (*Hoxd10 to Hoxd13)*. Mutations in this gene has been found in patients with Cornelia de Lange syndrome that have notably a clinodactyly of the 5th finger. Recently, a report showed that mice heterozygotes for *Nipbl* display polydactyly, and that lower dose of *Hoxd11* to *Hoxd13* in these mice further enhance this phenotype {Lopez-Burks, 2016 #45}. Other chromatin regulators were found enriched in this list of genes including *Jag1, Brd7, Jmjd6, Phf8, Ddb1, Hdac1, Swi5, Smarcc1, Smarce1, Hmgb3, Dnm3os, Cbx1* and *Lmnb1* (**Table S1**). The product of the latter gene has been associated with the architecture of a large domain of inactive chromatin (LADs, Kind, 2013 #25), where the *HoxD* cluster is not located (Vieux-Rochas et al PNAS 2015). Since reduced levels of *Lmnb1* gene product have been shown to be associated with reduced expression of polycomb targets including posterior *Hoxd* genes {Sadaie, 2013 #46}, its increased expression in the cells containing mRNAs from *Hoxd10* to *Hoxd13* may reflect a global change in chromatin configuration ({Reddy, 2008 #23};{Peric-Hupkes, 2010 #24};{Kind, 2013 #25};{Gonzalez-Sandoval, 2016 #26}). How would this change relate to a more permissive expression of *Hoxd* genes, as a cause or as a consequence, remains to be established.

The analyses of single cell transcriptomes revealed an unexpected hierarchical progression of *Hoxd* genes expression, from cells expressing a single posterior gene (*Hoxd13*) to the full combination (*Hoxd10 to Hoxd13)*. This global transcriptional sequence was inferred from a pseudo-time approach, a method whereby a temporal progression of cells is deduced based on their transcript patterns ({Haghverdi, 2016 #19};{Trapnell, 2014 #20}). We tested this hypothesis using diffusion pseudotime and found that autopod cells are much more subject to align along a developmental trajectory. This specificity may be link to the particular way *Hoxd* genes are regulated in distal limb buds, with a rapid and strong activation of *Hoxd13* due to its leading position in the cluster, towards the various enhancers ({Montavon, 2008 #4}). It is possible that the recruitment of additional *Hoxd* genes located nearby may be progressive, along with local epigenetic modifications, which could be inherited from one cell to its daughter cells. In this view, the number of *Hoxd* genes expressed would increase along with mitotic divisions leading to the hierarchical progression observed.

### Additive cellular or emerging functions?

The second question relates to the potential different functions that limb bud cells may display by carrying distinct combinations of *Hoxd* mRNAs. The question here is to discriminate between two views of the genotype-phenotype relationship during limb bud development, between a situation where each cell would express a determined combination of *Hoxd* mRNAs, for example in response to its topological position within the growing limb or to its own ‘regulatory history’, i.e. the regulations at work in its ancestor cells or, alternatively, a stochastic distribution of chromatin architectures leading to a globally balanced distribution of cells expressing various *Hoxd* mRNAs. In the former context, the resulting limb phenotype would derive from the additive effect of every single cells providing one out of the possible sets of information delivered by the various transcriptomes associated. In the second framework, the phenotype would derive from the random mixture of multiple cells expressing distinct transcriptomes with a given balance fixed by the choice of possible chromatin conformations.

Genetic approaches cannot easily discriminate between these alternatives. In previous studies where the functions of *Hoxd* genes during limb development were aimed to be assessed separately, various combinations of multiple inactivations were used. In most cases however, this consistently led to limited phenotypes due to a fair level of redundancy, particularly amongst *Hoxd* and *Hoxa* genes, preventing precise functions to be attributed to specific (groups of) *Hox* genes ({Zakany, 1997 #30}). However, the use of multiple gene inactivations has revealed that the transcription of *Hoxd11* and *Hoxd12* contributed functionally and thus added to the mere presence of *Hoxd13* transcripts, even though autopods double mutant for *Hoxd13* and *Hoxa13* would no longer grow ({Kmita, 2002 #21};{Delpretti, 2012 #17} {Fromental-Ramain, 1996 #28}). This is coherent with our data suggesting that the specific presence of *Hoxd11* or *Hoxd12* mRNAs is associated with distinct transcriptomes containing additional key regulators of cell fate and chromatin remodeling genes.

Therefore, part of the limb phenotypes observed in *Hoxd* multiple mutants may result from the different response of a sub group of cells, which would be differentially impacted by the loss of a given gene. For example, cells that express only *Hoxd13* or a combination of *Hoxd13* and *Hoxd10* mRNAs may be sensitive to the absence of *Hoxd11* transcripts in the corresponding mutant stock. Our results thus stress the necessity to keep in mind the cellular heterogeneity of transcriptional programs even in instances where WISH patterns seem to reveal homogenous distributions of transcripts. In this context, transcript patterns at the single-cell level can help solve the interpretation of genetically-deficient phenotypes, even though the co-regulation of *Hoxd* genes and the functional redundancy of their products make this statement difficult to apply to the present work.

Collectively, our observations revealed the existence of distinct combinations of *Hoxd* genes at the single-cell level during limb development. In addition, we document that the increasing combinatorial expression of *Hoxd* genes in this tissue is associated with specific transcriptional signatures and that these signatures illustrate a time progression in the differentiation of these cells. Further analysis at different developmental stages may enable the reconstruction of the cell fate trajectories and the state transitions that causes the cellular heterogeneity of the early limb bud tissue.

### METHODS

## Animal experimentation

All experiments were performed in agreement with the Swiss law on animal protection (LPA), under license No GE 81/14 (to DD). Forelimbs tissue samples were isolated from *Hoxd11::GFP* heterozygous animals at embryonic day 12.5 (E12.5) with day E0.5 being noon on the day of the vaginal plug. The cloning steps for the generation of the *Hoxd11* transgenic mice is described in **Fig. S1A-C**. Briefly, the knock-in was done by introducing a bi-cistronic cassette along with an IRES sequence. *Hoxd11* was inactivated by the insertion of a *TauGFP* sequence in frame into the coding sequence (**Fig. S1C**). The BamH1 site was used for insertion of the IRES cassette (**Fig. S1A**). **Fig. S1D** shows how the cassette was introduced as a single-copy knock-in. The GFP signal observed in this mouse stock reflects the endogenous distribution of *Hoxd11* transcription.

### RNA-FISH

E12.5 forelimbs were micro-dissected and fixed with 4% paraformaldehyde for 3 hours. Then the limbs were treated with sucrose at 5, 10 and 15% and then frozen in OCT. 25 μm cryostat sections were dried for 30 minutes, post-fixed in 4% paraformaldehyde for 10 minutes and quenched with 0.6% H2O2 in methanol for 20 minutes. Slides were then processed using the Ventana Discovery xT with the RiboMap kit. The pretreatment was performed with mild heating in CC2 for 12 minutes, followed by protease3 (Ventana, Roche) for 20 minutes at room temperature. Finally, the sections were hybridized using automated system (Ventana) with a *Hoxd13* probe diluted 1:1000 in ribohyde at 64°C for 6 hours. Three washes of 8 minutes in 2X SSC followed at hybridization temperature (64°C). Slides were incubated with anti-DIG POD (Roche Diagnostics) for 1 hour at 37°C in BSA 1% followed by a 10 minutes revelation with TSA substrate (Perkin Elmer) and 10 minutes DAPI. Slides were mounted in ProLong fluorogold. Images were acquired using a B/W CCD ORCA ER B7W Hamamatsu camera associated with an inverted Olympus IX81 microscope. The image stacks with a 2 μm step were saved as TIFF stacks. Image reconstruction and deconvolution were performed using FIJI (NIH, ImageJ v1.47q) and Huygens Remote Manager (Scientific Volume Imaging, version 3.0.3).

### RNA flow cytometry

Double *in situ* hybridization in single cells for RNA flow cytometry was performed using PrimeFlow RNA (Affymetrix, Santa Clara, CA) reagents following the manufacturer’s protocols. Cell viability was assessed by live/dead fixable dead cell; violet (ThermoFischer; L34955). Hsp90ab RNA probe (a gene expressed ubiquitously) served as a positive control. *Hoxd11* and *Hoxd13* RNA probes were used for the actual analysis. Cell staining was analyzed on a FACS Astrios located at the EPFL flow cytometry platform. Data analysis was performed by using FlowJoX (Treestar, Ashland, OR). The labelling and flow cytometry were performed on dissociated cells from eight forelimbs obtained from four different animals pooled together.

### Single-cell dissociation and fluorescence-activated cell sorting

Pools of embryonic forelimbs obtained from eight embryos were dissociated into a single-cell suspension using collagenase from Sigma (collagenase type XI) at 37C for 15min with 10 sec trituration. Cells were then resuspended in FACS solution (10% FCS in PBS with 2mM EDTA). Fluorescence activated cell sorting was performed using the MoFlow ASTRIOS EQ cell sorter with a 100-μ m nozzle. Through flow cytometry analysis performed using FlowJo (FlowJo LLC ©) we detected 1,602,844 cells positive for GFP in the autopod tissue and 235,000 simply negative. In the zeugopod tissue, 1,527,167 cells were positive, whereas 1,296,068 were negative giving thus a total of 87% GFP positive autopod cells and 54% positive zeugopod cells.

### Single-cell RNA sequencing, library preparation and mapping

Dissociated single-cells were obtained from eight *Hoxd11::GFP* forelimbs micro-dissected at E12.5. Cells with the highest level of GFP fluorescence (top 20%) were sorted using an Astrios cell sorter with a 100-μ m nozzle. 75bp large reads were uniquely mapped to the latest *Mus Musculus* reference genome (mm10) and the ERCC sequences using bowtie2 (REF:doi:10.1038/nmeth.1923) in local mode. Raw counts for the annotated ENSEMBL mouse genes (GRCm38) and the ERCC were obtained using the RNA-seq module of the HTSstation portal ({David, 2014 #274}). **Table S2** summarizes the raw counts. All single-cell RNA-seq data can be found in the Gene Expression Omnibus (GEO) repository under accession number GSEXX.

### Filtering low quality cells and genes expressed at low levels

Those counts were used to filter out some low-quality cells based on the following criteria: total number of reads mapped > 250, number of genes ‘expressed’ > 2000 (‘expressed’=with count > 0), and percent of reads mapped to Spike-Ins sequences < 25%. A total of 199 cells was retained (123 zeugopods and 76 autopods cells). Genes expressed at low levels were also removed from the rest of the analysis and only genes present (raw count > 0) in at least 10% of either the 76 autopods or the 123 zeugopods cells were retained. *Hox* genes were manually added if they did not satisfy these criteria. A total of 10’948 genes remained. ERCC with null counts through the remaining cells were also excluded from the rest of the analysis. **Table S4** summarizes those criteria and **Fig. S2**.

### Normalization

Raw counts were normalized with spike-in counts using the R package scran (methods used *computeSpikeFactors* and *normalize* version 1.0.4) (doi:10.18129/B9.bioc.scran). Prior to normalization, size factors were mean-centered to their batch of origin. An additional normalization step was also applied in order to correct for a potential gene length bias. **Table S5** compiles all the normalized values.

### Grouping of *Hoxd* gene combinations for differential gene expression analyses

*HoxD* groups were defined per cell and were composed by *Hoxd* genes with a minimum normalized expression of 5 when count represented at least 5% of the most expressed *Hoxd* genes in the cell. The differential gene expression analysis was performed with the R package *limma* (version 3.28.21) ({Ritchie, 2015 #42}). Genes with a minimum absolute log fold change of 2 and a BH adjusted p-value less than 0.01 (false discovery rate (FDR) of 1%) were considered differentially expressed.

### tSNE

The tSNE were computed using the package Rtsne (version 0.13) with the following parameters: 2 dimensions and a perplexity of 30, a maximum of iterations of 3000 and a seed set at 42. The top highly variable genes (HVG) that were used to plot the tSNE in **Fig. 3C-D** were selected using the trendVar & decomposeVar methods of the R pachage scran (version 1.0.4) (10.18129/B9.bioc.scran).

### Pseudo-time

Diffusion maps ({Coifman, 2005 #275}) are key tools to analyze single-cell differentiation data. It implements a distance metric relevant to how differentiation data is generated biologically, as cells follow noisy diffusion-like dynamics in the course of taking several differentiation lineage paths ({Haghverdi, 2016 #19}). The distances between cells reflect the transition probability based on several paths of random walks between the cells ({Angerer, 2016 #7}). The analysis was performed using the R package destiny (http://bioconductor.org/packages/destiny).

### Network visualization and Venn Diagram

Network shown in **Fig. 2** was built using weighted interaction networks from various sources of data and is able to process user data into such networks using a system that distinguishes between three different types of user-defined data in its import procedures: real- and binary-valued interaction networks, e.g. physical interaction networks; real-valued gene profile datasets, e.g. multi-sample microarray expression datasets; and binary-valued gene profile datasets {Mostafavi, 2008 #38}. Network shown in **Fig. 4** is a summary network of differentially expressed genes that was made with the R package Igraph (version 1.1.2; Csardi, G. & Nepusz, T. 2006)). The Igraph Software Package for Complex Network Research. InterJournal 2006, Complex Systems, 1695. http://igraph.org

## ACKNOWLEDGEMENTS

We thank B. Mangeat for his help with the Fluidigm C1 capture, as well as J. Faget and R. Colisson for their help with RNA prime flow labelling and flow cytometry analysis. We thank L. Beccari and other members of the Duboule laboratories for discussions. We also thank P. Schwalie and A. Necsulea for their help in the early steps of the single-cell RNA-seq analysis. Cell sorting was performed at the EPFL Flow Cytometry Core Facility. This work was supported by funds from the Ecole Polytechnique Fédérale (Lausanne), the University of Geneva, the Swiss National Research Fund (No. 310030B_138662) and the European Research Council grants System*Hox* (No 232790) and Regul*Hox* (No 588029)(to D.D.), as well as the SNF Ambizione grant PZ00P3_174032 (to PJF).

